# AI-guided Protein Inhibitor Design for Modulating FAD-dependent Glucose Dehydrogenase Redox Output

**DOI:** 10.64898/2026.07.20.739394

**Authors:** Su-Jin Lee, Eun-Ji Tak, Hyun-Jo Shim, Woo-Chan Ahn, Kwang-Hyun Park, Seong-Ryung Go, Haesik Yang, Eui-Jeon Woo

## Abstract

Flavin adenine dinucleotide-dependent glucose dehydrogenase (FAD-GDH) is a redox enzyme widely used in glucose monitoring, bioelectronic devices, and enzymatic biofuel cells because of its oxygen-independent catalysis and compatibility with electron-transfer processes. However, protein-based regulators that directly bind GDH and modulate its redox output remain underdeveloped. Here, we present an AI-guided strategy for developing a de novo protein inhibitor targeting FAD-GDH. GDH-targeting candidates generated through structure-based computational design were evaluated by yeast surface display and fluorescence-activated cell sorting, leading to the identification of FAD-GDH inhibitor-1 (FGI-1) as a GDH-targeting inhibitory scaffold. Purified His–MBP–FGI-1 reduced GDH-mediated DCIP reduction, demonstrating attenuation of GDH-derived redox output. Random mutagenesis followed by secondary FACS screening yielded evolved variants with increased GDH-binding signals and enhanced redox-output suppression, showing that the de novo inhibitory scaffold could be functionally tuned through experimental evolution. In addition, an FGI-1-based construct fused to a larger protein module retained GDH-output suppressive activity, and electrode-based measurements showed reduced GDH-derived current output. Because electrode-associated measurements may be influenced by protein-mediated surface shielding and altered electron-transfer accessibility, this decrease was interpreted conservatively as attenuation of GDH-derived electrochemical output rather than direct evidence of active-site inhibition. Together, this work establishes an AI-guided design–validation workflow for developing protein inhibitors that modulate FAD-GDH redox output and provides a foundation for protein-level control of enzyme output in biosensing and bioelectronic applications.

## Introduction

Glucose dehydrogenase (GDH) is a glucose-responsive redox enzyme that converts glucose oxidation into measurable electron-transfer output and has been widely applied in glucose monitoring, biosensing, bioelectronic devices, and enzymatic biofuel cells **[1–4]**. Among the different GDH classes, flavin adenine dinucleotide-dependent glucose dehydrogenase (FAD-GDH) is particularly attractive because it contains a tightly bound FAD cofactor and catalyzes glucose oxidation independently of molecular oxygen **[2,4]**. Unlike glucose oxidase, which uses oxygen as an electron acceptor and generates hydrogen peroxide, FAD-GDH supports oxygen-insensitive redox cycling without producing hydrogen peroxide, making it well suited for sensing and bioelectronic applications under variable oxygen conditions **[2,4]**.

Substrate specificity varies among fungal FAD-GDHs, although several enzymes developed for glucose sensing exhibit favorable selectivity toward glucose **[2,4–6]**. This property is important for generating glucose-dependent signals in biological samples containing multiple saccharides. In addition, FAD-GDH has been investigated as an electrode-compatible biocatalyst capable of supporting direct or mediated electron transfer **[3,7]**. These properties make FAD-GDH a useful redox module that combines oxygen-independent catalysis, glucose recognition, and compatibility with electrochemical signal generation.

In conventional GDH-based systems, redox output has been optimized by adjusting enzyme loading, immobilization matrices, mediator composition, electrode architecture, and reaction conditions **[1,7,8]**. Although these engineering approaches have substantially improved device performance, they mainly regulate the environment surrounding the enzyme rather than directly controlling GDH through molecular recognition. A protein inhibitor that binds GDH and attenuates its redox output could therefore provide a complementary strategy for controlling enzyme function without changing the amount of enzyme or the overall device architecture.

Despite the broad utility of FAD-GDH, protein-based regulators that directly modulate its redox output remain poorly developed. Previous studies have focused primarily on enzyme discovery, biochemical characterization, stabilization, structural analysis, and engineering of substrate specificity or electron-transfer properties **[4–6,9,10]**. To our knowledge, a de novo designed protein inhibitor capable of suppressing FAD-GDH-mediated redox output has not been reported. Such an inhibitor could serve as a molecular tool for probing GDH function and provide a new approach for tuning enzyme-derived signals in biosensing and bioelectronic systems.

Recent advances in AI-guided protein design have enabled the generation of synthetic binders and functional protein modulators that are not derived from naturally occurring interaction partners **[11–13]**. Generative backbone design, sequence optimization, structure prediction, and display-based screening can now be integrated to design and experimentally identify binders against selected protein surfaces **[14–18]**. Directed evolution and targeted mutagenesis can subsequently refine initially identified scaffolds by improving their folding, target engagement, or functional activity **[19]**. Applying this combined computational and experimental strategy to redox enzymes offers a route to develop artificial regulatory proteins that control enzyme output through designed surface recognition.

Here, we report an AI-guided design and validation workflow for developing a de novo protein inhibitor of FAD-GDH. GDH-targeting helical scaffolds were computationally generated, screened by yeast surface display and fluorescence-activated cell sorting, and functionally evaluated using GDH-mediated redox assays. The selected inhibitor, FGI-1, attenuated GDH-catalyzed DCIP reduction. Random mutagenesis and secondary FACS screening subsequently yielded evolved variants with increased GDH-binding signals and enhanced redox-output suppression. An FGI-1-based fusion construct also retained suppressive activity when incorporated into a larger protein architecture and reduced GDH-derived output in an electrode-based assay. Together, these results establish an AI-guided design–validation framework for developing protein regulators of FAD-GDH and provide a foundation for protein-based control of enzyme-derived redox output in biosensing and bioelectronic applications

## Results and discussion

### AI-guided design and identification of FGI-1

To develop a protein-based regulator of flavin adenine dinucleotide-dependent glucose dehydrogenase (FAD-GDH), we established an integrated computational and experimental workflow for designing de novo binders capable of attenuating GDH-derived redox output. FAD-GDH catalyzes glucose oxidation through FAD-dependent electron transfer, and the resulting electron flow can be coupled to an electrode-based signal **(Figure 1A)**. We therefore reasoned that a compact protein binder positioned near the catalytic entrance or an adjacent surface region could attenuate GDH output by restricting productive substrate access or altering local redox coupling.

**Figure 1.**
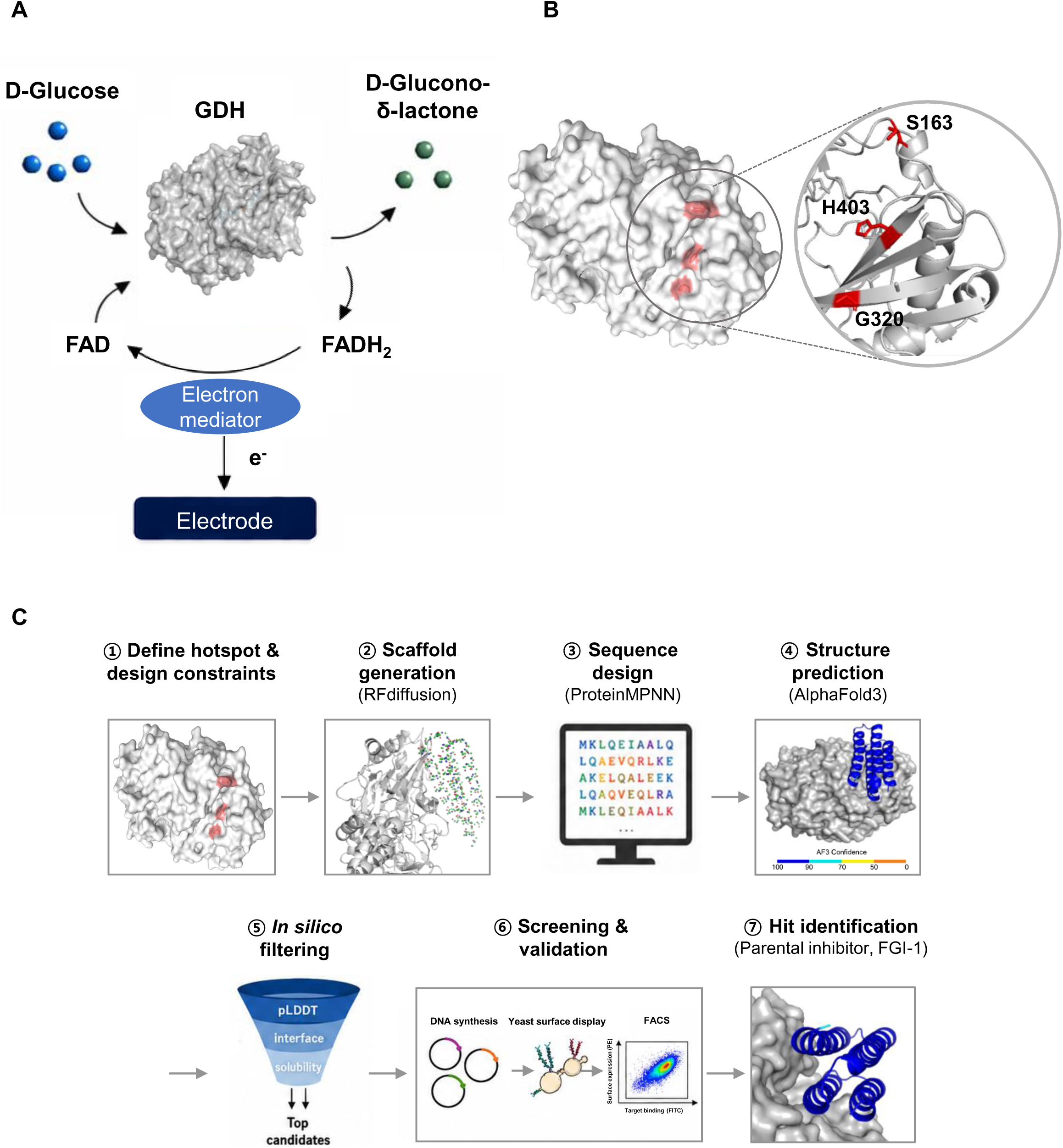
AI-guided design of a de novo FAD-GDH inhibitor. **(A)** Schematic of FAD-GDH-catalyzed glucose oxidation and electron transfer. **(B)** FAD-GDH surface showing the selected hotspot residues S163, G320, and H403. **(C)** Computational design and experimental screening workflow, including RFdiffusion, ProteinMPNN, AlphaFold3, in silico filtering, yeast surface display, and FACS, leading to identification of the parental inhibitor FGI-1.

The crystal structure of *Aspergillus flavus* FAD-GDH (PDB:4YNU) was used as the structural basis for target preparation **[10]**. Surface-exposed residues S163, G320, and H403, located near the catalytic loop and FAD-adjacent region, were selected as candidate hotspots for binder generation **(Figure 1B)**.

De novo binder backbones were generated using RFdiffusion, with the selected GDH surface region specified as the design target **[14]**. A compact four-helix bundle scaffold was selected based on predicted surface complementarity and overall structural compactness. The resulting scaffold was subsequently modeled against WT FAD-GDH and subjected to sequence optimization using ProteinMPNN **[15]**. Candidate sequences were generated across multiple sampling temperatures and filtered according to predicted solubility, monomeric folding confidence, and complex-interface quality. AlphaFold3 prediction of the resulting complex was used to confirm the binder geometry against WT GDH model **[17] (Figure S1)**. Overall, 500 sequences were generated, of which 88 passed the computational filters. Six candidates were initially prioritized by visual inspection, while the remaining 82 were subsequently evaluated in the same screening framework **(Table S1)**.

Selected candidates were displayed on the surface of *Saccharomyces cerevisiae* and analyzed by fluorescence-activated cell sorting. Surface expression was monitored using an anti-Myc antibody, whereas binding to His-tagged GDH was detected using an anti-His fluorescent signal. Among the six initially prioritized candidates, one clone showed a reproducible dual-positive population and was enriched during sorting. Screening of the remaining 82 candidates did not yield an additional detectable binder under the conditions used. Sequence analysis of the enriched population identified a dominant clone, which was designated FAD-GDH inhibitor-1 (FGI-1) and selected for subsequent functional and structural characterization. Together, these results establish an AI-guided design-to-screening workflow that combines generative backbone design, sequence optimization, structure prediction, and yeast surface display to identify a de novo GDH-targeting protein scaffold **(Figure 1C)**.

### Functional and structural characterization of FGI-1

To further evaluate the GDH-binding properties of FGI-1, iterative sorting was performed using yeast surface display. The fraction of cells positive for both surface expression and GDH binding was initially low but progressively increased over three rounds of sorting **(Figure 2A)**. This enrichment indicates that FGI-1 reproducibly engages GDH when displayed on the yeast surface.

**Figure 2.**
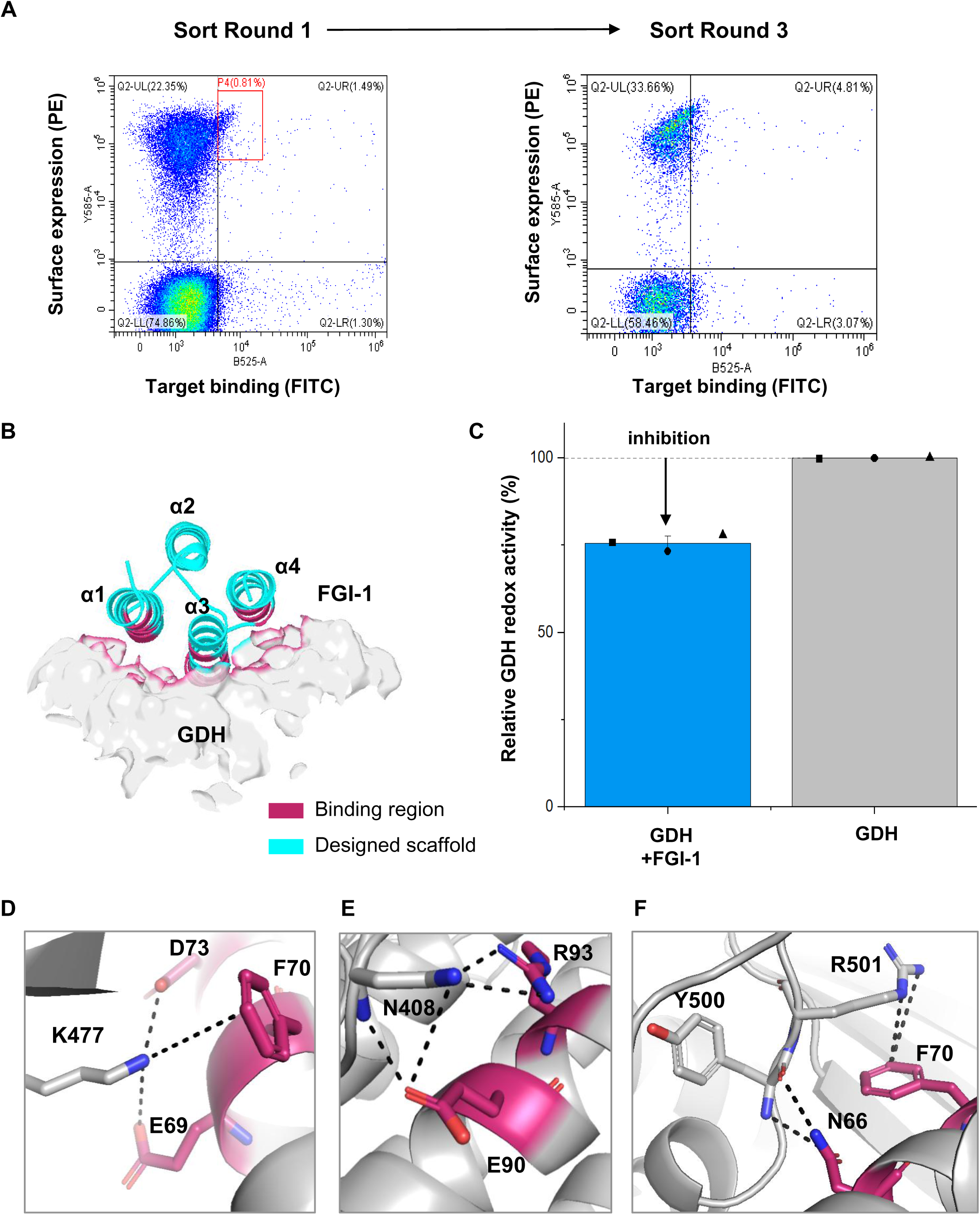
Identification and structural characterization of FGI-1. **(A)** Representative FACS plots showing enrichment of GDH-binding yeast during sorting. **(B)** Predicted GDH–FGI-1 complex, with the binding region highlighted in magenta and the remaining scaffold in cyan. **(C)** Relative GDH redox activity in the presence or absence of FGI-1. **(D–F)** Predicted interface interactions involving GDH residues K477, N408, Y500, and R501. Data are shown as mean ± SD with individual measurements.

To examine the predicted binding mode of FGI-1, we analyzed the AlphaFold3-predicted GDH– FGI-1 complex **(Figure 2B)**. In the model, FGI-1 adopts a compact four-helix scaffold and binds to a surface region adjacent to the catalytic entrance of GDH. A subset of FGI-1 residues forms the predicted GDH-contacting interface, whereas the remaining scaffold supports the overall fold. This binding geometry suggests that FGI-1 may influence GDH function by occupying a surface near the catalytic entrance rather than inserting deeply into the enzyme core, potentially affecting productive substrate access or local redox coupling.

FGI-1 was expressed and purified as an N-terminal His–MBP fusion protein, and the purified protein was used for the functional assay **(Figure S2)**. We next assessed whether FGI-1 could modulate GDH-derived redox output using purified His–MBP–FGI-1 in a DCIP-based assay. When the activity of GDH alone was normalized to 100%, addition of FGI-1 reduced the relative GDH redox activity to approximately 75% **(Figure 2C)**. These results indicate that FGI-1 not only binds GDH but also attenuates GDH-mediated electron-transfer output. Because DCIP reduction reflects the overall GDH-coupled redox reaction, including enzymatic glucose oxidation and subsequent electron transfer to the redox mediator, the observed decrease indicates reduced apparent GDH redox activity. However, without detailed kinetic analysis, this assay does not distinguish whether the effect arises from changes in intrinsic catalytic turnover, substrate accessibility, or electron-transfer efficiency.

The predicted GDH–FGI-1 interface contained several polar and electrostatic interaction clusters. GDH residue K477 was predicted to interact electrostatically with FGI-1 residues E69 and D73 and to form a cation–π contact with F70 **(Figure 2D)**. GDH residue N408 was positioned within a polar interaction network involving FGI-1 residues E90 and R93 **(Figure 2E)**. Additional anchoring contacts were predicted around GDH residues Y500 and R501, involving FGI-1 residues N66 and F70 **(Figure 2F)**. Although these contacts are based on a computational model, they provide a plausible structural basis for the observed GDH-binding and redox-output suppressive activity of FGI-1.

Together, these results show that FGI-1 was enriched for GDH binding during iterative yeast surface display sorting and attenuated GDH-mediated DCIP reduction in the purified His–MBP fusion format. The predicted complex suggests that this functional effect may arise from multivalent polar and electrostatic interactions formed at a surface region adjacent to the catalytic entrance.

### Directed evolution of FGI-1 enhances GDH-binding signals and redox-output suppression

To determine whether the GDH-targeting properties of FGI-1 could be further improved, we subjected the parental scaffold to directed evolution. An error-prone PCR library was generated from the FGI-1 sequence, displayed on the surface of yeast, and screened by FACS using GDH as the target **(Figure 3A)**. Variants exhibiting increased target-binding signals while maintaining detectable surface expression were enriched and recovered for sequence analysis.

**Figure 3.**
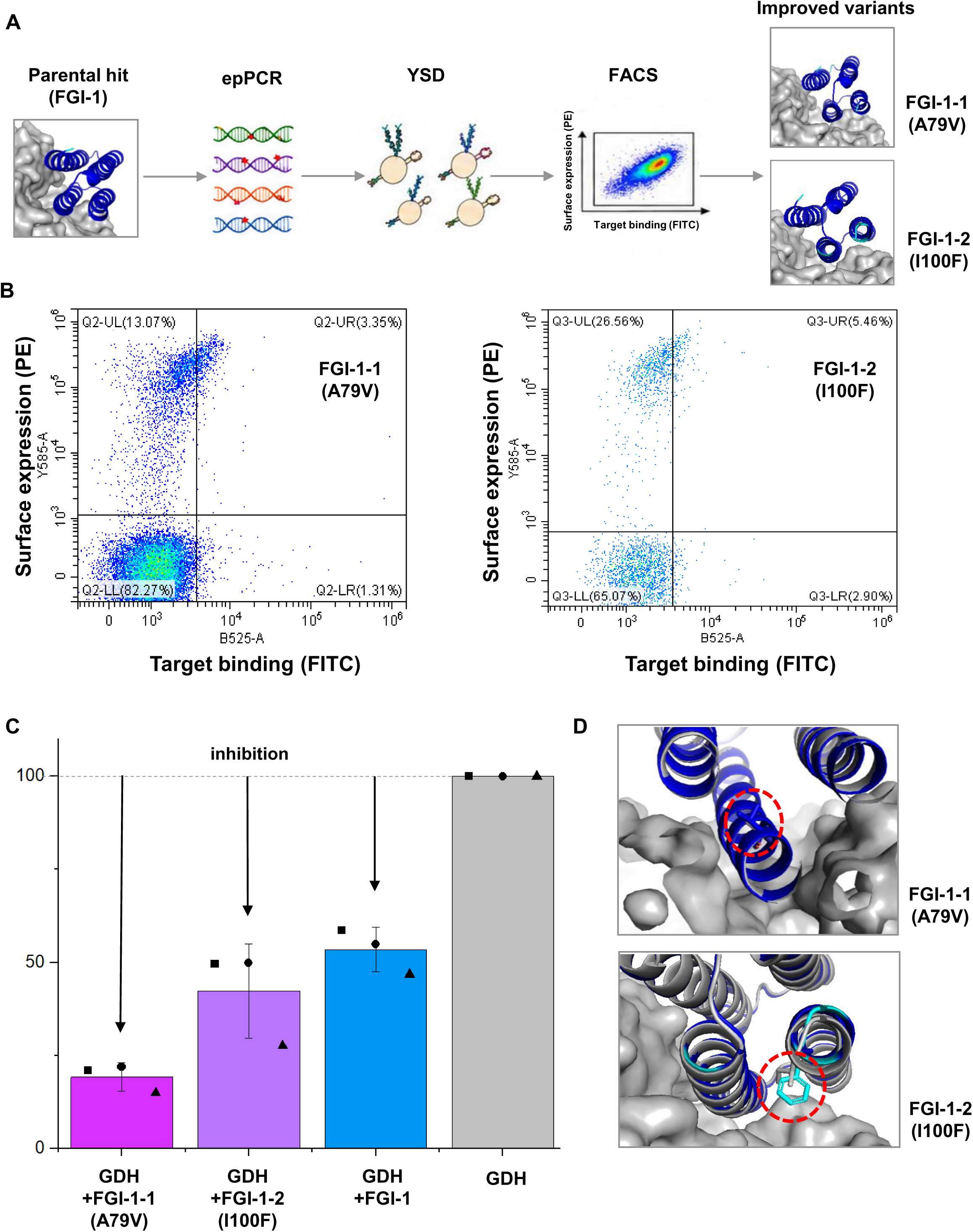
Directed evolution of FGI-1. Error-prone PCR, yeast surface display, and FACS workflow used to identify improved variants. **(A)** Representative FACS plots of FGI-1-1 (A79V) and FGI-1-2 (I100F). **(C)** Relative GDH redox activity in the presence of parental and evolved inhibitors. **(D)** Predicted structural locations of A79V and I100F. Data are shown as mean ± SD with individual measurements.

Compared with parental FGI-1, the enriched variants showed increased GDH-binding populations in the yeast surface display assay. Two representative variants carrying the substitutions A79V and I100F were selected and designated FGI-1-1 and FGI-1-2, respectively **(Figure 3B)**. Because the FACS readout reflects both surface presentation and target engagement, the increased fluorescence signals were interpreted as improved apparent GDH-binding behavior under the yeast display conditions rather than as direct measurements of binding affinity.

To assess whether the increased binding signals were accompanied by improved functional activity, FGI-1-1 and FGI-1-2 were similarly purified as N-terminal His–MBP fusion proteins **(Figure S2)** and evaluated using the DCIP-based GDH redox assay. Both evolved variants produced greater attenuation of GDH-mediated redox activity than the parental FGI-1 under matched assay conditions **(Figure 3C)**. These results indicate that sequence diversification and FACS-based enrichment can enhance the redox-output suppressive function of the de novo GDH-targeting scaffold.

Structural models were subsequently examined to explore the potential effects of the selected substitutions **(Figure 3D)**. A79 is located within the helical scaffold, and its substitution with valine may alter local hydrophobic packing or scaffold stability. I100F introduces a larger aromatic side chain at another position within the binder and may influence local packing or the presentation of the GDH-contacting surface. However, because these interpretations are based on predicted structures, the precise molecular contributions of A79V and I100F to GDH recognition and functional suppression remain to be experimentally established. Together, these results demonstrate that FGI-1 is an evolvable inhibitory scaffold. Directed evolution increased GDH-binding signals in the yeast display assay and yielded variants with enhanced suppression of GDH-derived redox output.

### Functional retention of an FGI-1-based modular fusion and attenuation of electrochemical GDH output

To determine whether FGI-1 could retain its GDH-output suppressive function when incorporated into a larger protein architecture, FGI-1 was genetically connected to a large protein module through a flexible linker **(Figure 4A)**. The construct was expressed as an N-terminal His–MBP fusion. The FGI-1–large-module fusion protein was purified after TEV-mediated removal of the His–MBP tag **(Figure S2).** The resulting fusion protein was then used for subsequent functional analyses.

**Figure 4.**
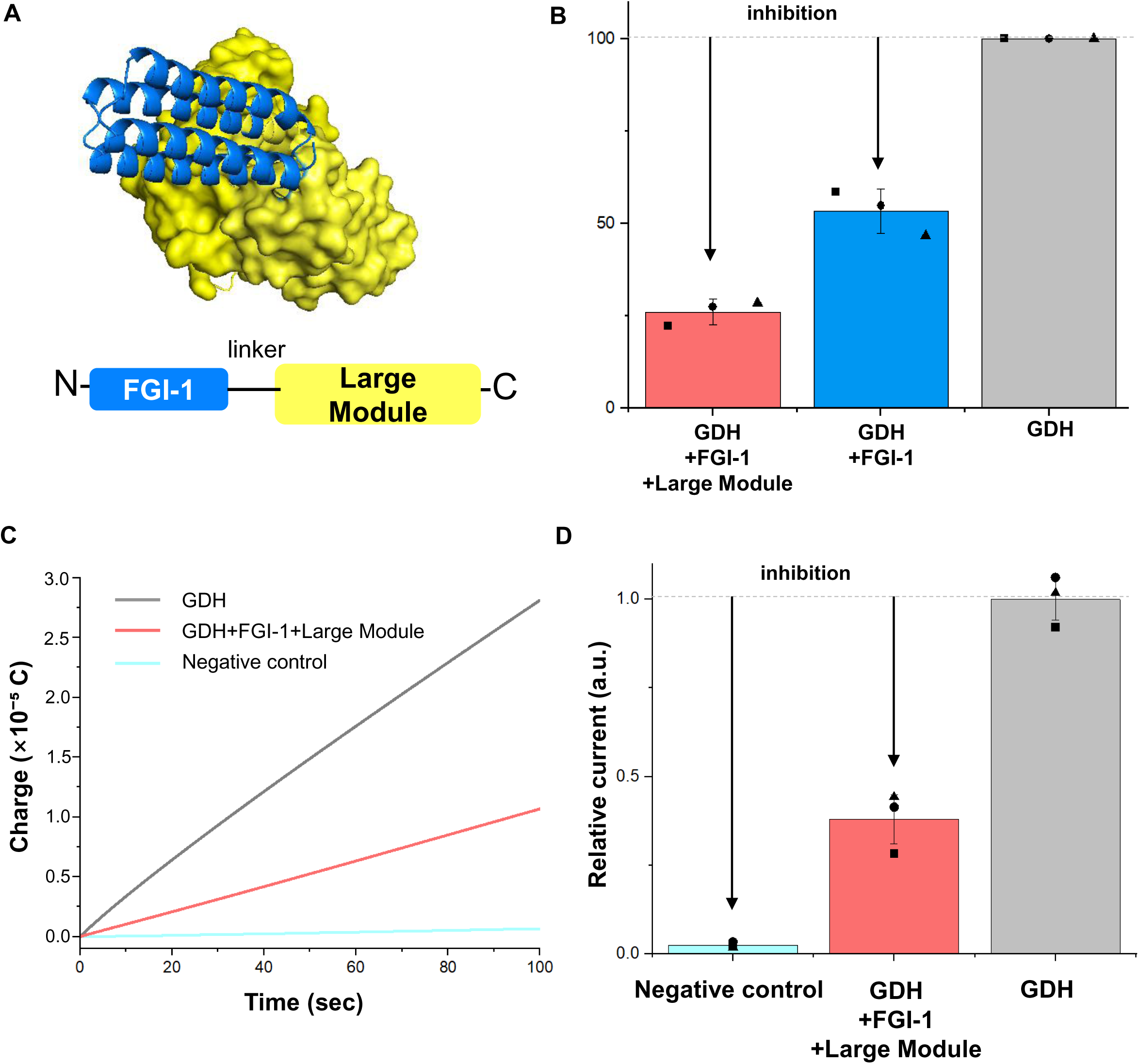
Functional evaluation of an FGI-1 modular fusion. **(A)** Structural model and schematic of FGI-1 fused to a large protein module. **(B)** Relative GDH redox activity in the presence of FGI-1 or the fusion construct. **(C, D)** Representative electrochemical charge traces and relative current output for the negative control, GDH alone, and GDH with the fusion construct. Data are shown as mean ± SD with individual measurements.

We first assessed the ability of the modular fusion protein to attenuate GDH-derived redox output using the DCIP-based assay. Relative to the GDH-only condition, both FGI-1 and the FGI-1–large-module fusion protein reduced GDH-mediated redox activity **(Figure 4B)**. The retention of suppressive activity after fusion to a substantially larger protein module indicates that FGI-1 can function as a modular inhibitory element and remains compatible with an expanded protein architecture.

We next investigated whether the FGI-1-based fusion protein could also reduce GDH-derived output in an electrode-associated measurement. GDH and the FGI-1–large-module fusion protein were mixed in solution and applied together to the electrode, after which accumulated charge was monitored over time **(Figure 4C, D)**. GDH alone produced a clear time-dependent increase in accumulated charge, whereas the condition containing both GDH and the fusion protein showed reduced charge accumulation and lower relative current output. These results suggest that the FGI-1-based fusion protein attenuates GDH-derived electrochemical output in addition to suppressing the solution-phase DCIP response.

However, the decrease in current observed in the electrode-based assay does not necessarily indicate direct inhibition of the GDH active site. Binding of the FGI-1–large-module fusion protein may sterically restrict access of the substrate or redox mediator or alter the accessibility of electron-transfer pathways from GDH to the electrode. Therefore, these results were interpreted as showing that the FGI-1-based fusion protein attenuates GDH-derived electrochemical output through direct or indirect inhibitory effects. Together, these results demonstrate that FGI-1 retains its GDH-output suppressive function after fusion to a larger protein module and reduces GDH-derived current output in an electrode-based assay. This functional compatibility supports the potential use of FGI-1 as a portable inhibitory element that can be incorporated into larger protein or sensor-oriented architectures.

## Methods

### Expression and purification of WT FAD-GDH

The gene encoding flavin adenine dinucleotide-dependent glucose dehydrogenase (FAD-GDH) from *Aspergillus flavus* was cloned into the pET-30a(+) vector containing a C-terminal His₆ tag. The recombinant plasmid was transformed into *Escherichia coli* BL21(DE3), and protein expression was induced with 0.5 mM isopropyl β-D-1-thiogalactopyranoside (IPTG) at 18 °C for 18 h. Cells were harvested and resuspended in 10 mM MOPS buffer (pH 6.5) containing 5% glycerol and 5 mM β-mercaptoethanol. The cells were disrupted by sonication, and cell debris was removed by centrifugation at 13,000 × *g* for 40 min. The supernatant was purified sequentially by Q anion-exchange chromatography and Ni-NTA affinity chromatography. Purified FAD-GDH was analyzed by SDS–PAGE, dialyzed against 10 mM MOPS buffer (pH 6.5) containing 5% glycerol and 5 mM β-mercaptoethanol, aliquoted, and stored at −80 °C.

### AI-guided design of FAD-GDH-binding proteins

The crystal structure of *A. flavus* FAD-GDH (PDB ID: 4YNU) was used as structural template **[10, 17]**. Residues S163, G320, and H403 were designated as hotspots to guide binder-interface formation. De novo binder backbones were generated using RFdiffusion **[14]** in the ColabDesign v1.1.1 environment. Residues 1–571 of GDH chain A were defined as the target contig, and α-helical binder scaffolds of 100–150 amino acids were generated. Fifty diffusion trajectories were generated for WT GDH template, yielding a total of 50 initial backbone models. Among the structure generated using the WT GDH template, a 107-residue four-helix bundle scaffold with favorable predicted interface placement and structural compactness was selected, and the WT GDH–binder complex with an improved predicted interface was used as the template for subsequent sequence design. Binder sequences were optimized using ProteinMPNN **[15]**. Sampling temperatures of 0.05, 0.10, 0.15, 0.20, and 0.25 were used, and 100 sequences were generated at each temperature, resulting in a total of 500 candidate sequences. Predicted solubility was assessed using SoluProt, and candidates with SoluProt scores of at least 0.7 were prioritized. GDH–binder complexes were predicted using AlphaFold3 **[17]**. Candidates were evaluated based on binder folding confidence, complex-structure confidence, continuity of the predicted interface, surface complementarity, polar and electrostatic interactions, and aromatic or cation–π contacts. A total of 88 candidates passed the computational filters. Six candidates were prioritized for initial experimental validation, and the remaining 82 candidates were evaluated using the same yeast surface display-based screening workflow.

### Yeast surface display and FACS-based binder screening

Genes encoding the designed binders were synthesized as gBlock or eBlock DNA fragments (Integrated DNA Technologies). Each gene was cloned into the pCTCON3 yeast surface display vector for expression as an Aga2 fusion protein and introduced into *Saccharomyces cerevisiae* EBY100 by electroporation **[18]**. Transformed yeast cells were expanded in SDCAA medium and induced in SGCAA medium to express the binders on the cell surface. Following induction, cells were washed with phosphate-buffered saline containing 1% bovine serum albumin (PBSA). Binder surface expression was detected using a PE-conjugated anti-Myc antibody, whereas GDH binding was detected using His-tagged FAD-GDH and an anti-His FITC antibody. For initial screening, 1 µM His-tagged FAD-GDH was preincubated with anti-His FITC on ice for 15 min before being added to the yeast cells. Anti-Myc PE was then added, and the cells were stained at 4 °C for 1 h. FITC and PE fluorescence channels were used to monitor GDH binding and binder surface expression, respectively. Cells positive for both fluorescence signals were isolated by fluorescence-activated cell sorting. Sorted cells were recovered in SDCAA medium, and iterative rounds of sorting were performed using the same procedure. The binding ratio was calculated as the fraction of FITC/PE double-positive cells within the total gated yeast population. Plasmids were recovered from sorted yeast cells and transformed into *E. coli* DH10B. Individual colonies were analyzed by Sanger sequencing. A dominant GDH-binding clone repeatedly identified among the recovered sequences was designated FAD-GDH inhibitor-1 (FGI-1).

### Expression and purification of FGI-1

The FGI-1 gene was cloned into the pMAL-c2X vector downstream of an N-terminal His₁₀–MBP fusion tag. The recombinant plasmid was transformed into *E. coli* BL21(DE3), and expression was induced with 0.5 mM IPTG at 18 °C for 18 h. Cells were harvested and resuspended in buffer A containing 300 mM NaCl, 30 mM Tris-HCl (pH 8.0), 10% glycerol, and 5 mM β-mercaptoethanol. The cells were disrupted by sonication and centrifuged at 13,000 × *g* for 40 min. The supernatant was applied to Ni-NTA affinity resin to purify the His₁₀–MBP–FGI-1 fusion protein. The fusion protein was further purified by Q anion-exchange chromatography and Superdex 75 size-exclusion chromatography. Purified His₁₀–MBP–FGI-1 was dialyzed against PBS before functional analysis. Final protein purity and apparent molecular weight were assessed by SDS–PAGE, and purified His₁₀– MBP–FGI-1 was used in the functional assay shown in **Figure 2C**.

### Expression and purification of evolved FGI-1 variants

The FGI-1-1 and FGI-1-2 genes were cloned into the pMAL-c2X vector downstream of an N-terminal His₁₀–MBP fusion tag. Recombinant constructs were expressed in *E. coli* BL21(DE3). Protein expression was induced with 0.5 mM IPTG at 18 °C for 18 h. Cells were harvested and resuspended in PBS containing 10% glycerol, 5 mM β-mercaptoethanol, and a protease inhibitor cocktail, disrupted by sonication, and centrifuged at 13,000 × *g* for 40 min. Clarified lysates were incubated with Ni-NTA affinity beads to allow batch binding of the His₁₀–MBP-tagged proteins. The protein-bound beads were subsequently transferred to an open gravity-flow column, washed with the binding buffer, and eluted with PBS containing 10% glycerol, 5 mM β-mercaptoethanol, and 250 mM imidazole. The evolved variants were retained as His–MBP fusion proteins and dialyzed against PBS before functional analysis. They were then used in the functional assays shown in **Figure 3**. Protein purity and apparent molecular weight were assessed by SDS–PAGE.

### Construction and purification of the FGI-1–large-module fusion protein

FGI-1 was genetically connected to a large protein module through a flexible peptide linker. The fusion construct was expressed in *E. coli* BL21(DE3) as an N-terminal His–MBP fusion protein and purified by Ni-NTA affinity chromatography. The purified fusion protein was incubated with TEV protease at 4 °C to remove the His–MBP tag. The sample was subsequently reapplied to Ni-NTA resin to remove the cleaved His–MBP tag, His-tagged TEV protease, and uncleaved fusion protein. The flow-through containing the tag-cleaved FGI-1–large-module fusion protein was collected. The purified fusion protein was analyzed by SDS–PAGE and used for the DCIP-based assay and electrochemical measurements shown in **Figure 4**.

### DCIP-based analysis of redox-output suppression by FGI-1

The activity of FGI-1 shown in **Figure 2C** was evaluated using FGI-1 in a DCIP-based assay **[2]**. Each 120 µL reaction contained WT FAD-GDH and FGI-1 at final concentrations of 7.8 µM and 65.04 µM, respectively. GDH and FGI-1 were incubated at 25 °C for 15 min. DCIP was used as an artificial electron acceptor, and the decrease in absorbance at 600 nm resulting from DCIP reduction during glucose oxidation was measured. The redox activity of the GDH-only condition was normalized to 100%, and the relative activity of the FGI-1-containing condition was calculated. All experiments were performed in triplicate, and data are presented as mean ± standard deviation.

### DCIP-based analysis of evolved variants and the large-module fusion protein

The functional assays shown in **Figures 3 and 4** were performed in a total reaction volume of 200 µL. Each reaction contained 160 µL of a 10 mM phosphate reaction mixture (pH 6.5) containing DCIP, 20 µL of 400 mM glucose, 5 µL of 0.052 µM GDH stock, 10 µL of 1.52 µM inhibitor stock, and 5 µL of PBS. The final concentrations of glucose, GDH, and inhibitor were therefore 40 mM, 1.3 nM, and 0.076 µM, respectively. His–MBP–FGI-1, His–MBP–FGI-1-1, or His–MBP–FGI-1-2 was used as the inhibitor in the assays shown in **Figure 3**. The tag-cleaved FGI-1–large-module fusion protein was used at the same final concentration in the assay shown in **Figure 4**. For the GDH-only control, the inhibitor volume was replaced with an equal volume of PBS to maintain the same total reaction volume and buffer composition. Reactions were initiated by the addition of glucose, and changes in absorbance at 600 nm associated with DCIP reduction were monitored. The redox activity of the GDH-only condition was normalized to 100%, and the relative activities of the inhibitor-containing conditions were calculated. Each condition was measured in triplicate, and data are presented as mean ± standard deviation.

### Error-prone PCR-based directed evolution of FGI-1

The FGI-1 gene was used as the template for error-prone PCR to generate mutant libraries. Two mutagenic PCR conditions were used to obtain different mutation rates, corresponding to an average of approximately two or five substitutions per gene. Amplified DNA fragments were purified and cloned into the pCTCON3 yeast surface display vector. The assembled libraries were introduced into *S. cerevisiae* EBY100 by electroporation. Following induction of surface expression, the libraries were stained using anti-Myc PE and His-tagged FAD-GDH pre-complexed with anti-His FITC. Cells that retained binder surface expression while exhibiting higher GDH-binding fluorescence than parental FGI-1 were isolated by FACS. Sorted populations were recovered in SDCAA medium, and the recovered plasmids were transformed into *E. coli* DH10B. Individual colonies were analyzed by Sanger sequencing. Representative evolved variants containing the A79V and I100F substitutions were designated FGI-1-1 and FGI-1-2, respectively. These proteins were expressed and purified as N-terminal His–MBP fusion proteins and evaluated using the DCIP-based GDH redox assay.

### Structural modeling and interface analysis

The GDH–FGI-1 complex structure was predicted using AlphaFold3 **[17]**. The predicted model was analyzed to examine the overall fold of FGI-1, its binding position on the GDH surface, and residue-level contacts at the GDH–FGI-1, FGI-1-1 and FGI-1-2 interface. All interpretations of interface contacts and mutation effects were considered model-based predictions. Structural visualization and figure preparation were performed using PyMOL v3.0, and protein–protein interfaces were analyzed using PDBePISA.

### Preparation of ITO electrodes containing the FGI-1–large-module fusion protein

Indium tin oxide (ITO) electrodes were used for electrochemical analysis. For preparation of electrodes containing the FGI-1–large-module fusion protein, 5 µL of 10 µg/mL WT GDH stock, 5 µL of 9.2 µM FGI-1–large-module fusion protein, and 40 µL of PBS were mixed to a total volume of 50 µL. The GDH-only control was prepared by mixing 5 µL of 10 µg/mL WT GDH stock with 45 µL of PBS to the same total volume. Each mixture was incubated at 25 °C for 15 min and then drop-cast onto an ITO electrode in a volume of 10 µL. The electrodes were dried at 25 °C for 30 min with the incubator lid open to allow solvent evaporation. The dried ITO electrodes containing adsorbed GDH alone or GDH together with the FGI-1–large-module fusion protein was used for electrochemical measurements.

### Electrochemical measurement of GDH-derived output

1,10-Phenanthroline-5,6-dione (PD) was used as a redox mediator. The glucose-containing measurement solution was prepared by mixing 100 µL of 50 mM glucose, 100 µL of 1 mM PD, and 800 µL of PBS (pH 7.4) to a final volume of 1 mL. The final concentrations of glucose and PD were 5 mM and 0.1 mM, respectively. The background solution was prepared by mixing 100 µL of 1 mM PD with 900 µL of PBS to a final volume of 1 mL. Prepared electrodes were immersed in the corresponding measurement solutions, and current signals were recorded using a CH Instruments electrochemical workstation. Chronocoulometry was performed at an applied potential of 0.0 V in PBS buffer (pH 7.4). The following signals were compared: the background signal measured in PD/PBS solution without glucose, the glucose-dependent signal from electrodes prepared by co-deposition of GDH and the FGI-1–large-module fusion protein, and the glucose-dependent signal from GDH-only control electrodes prepared with PBS in place of the inhibitor. Current was integrated over time to calculate accumulated charge over 100 s. The FGI-1–large-module fusion protein condition was measured in two independent experiments.

### Statistical and data analysis

Quantitative data are presented as mean ± standard deviation. DCIP-based functional assays were performed in triplicate for each condition, whereas the electrochemical analysis was performed using three independent measurements. Relative GDH redox activity was calculated by normalizing the GDH-only control to 100%. The same gating strategy was applied to all flow-cytometry samples, and cells positive for both binder surface expression and GDH-binding fluorescence were defined as the double-positive population. For electrochemical analysis, current was integrated over time to calculate accumulated charge. Individual replicate values were displayed as points, bars represent the mean, and error bars indicate the standard deviation.

## Acknowledgements

This work was supported by National Research Foundation of Korea (NRF) grants funded by the Korean Government (MSIP) (RS-2022-NR071772, RS-2021-NR059435 and RS-2021-NR056526), the National Research Council of Science & Technology (NST) (CRC22024-500) and the KRIBB Research Initiative Program (KGM5382632, KGM1062612, KGM1322612).

## Author contributions

**Su-Jin Lee**: Conceptualization, Methodology, Software, Validation, Formal analysis, Investigation, Data curation, Visualization, Writing – original draft, Writing – review & editing. **Eun-Ji Tak**: Methodology, Validation, Investigation, Formal analysis, Data curation. **Hyun-Jo Shim**: Investigation, Formal analysis, Data curation. **Woo-Chan Ahn**: Software, Investigation, Formal analysis, Data curation. **Kwang-Hyun Park**: Software, Investigation, Formal analysis, Data curation. **Seong-Ryung Go**: Investigation, Formal analysis, Data curation. **Haesik Yang**: Methodology, Resources, Validation, Investigation, Formal analysis, Data curation. **Eui-Jeon Woo**: Conceptualization, Supervision, Project administration, Funding acquisition, Writing – review & editing. All authors have read and approved the final version of the manuscript.

## Competing interests

A patent application related to the FGI-1-based FAD-GDH inhibitor platform described in this manuscript has been filed by the authors and/or their institutions.

## Supporting information

**Figure S1.**
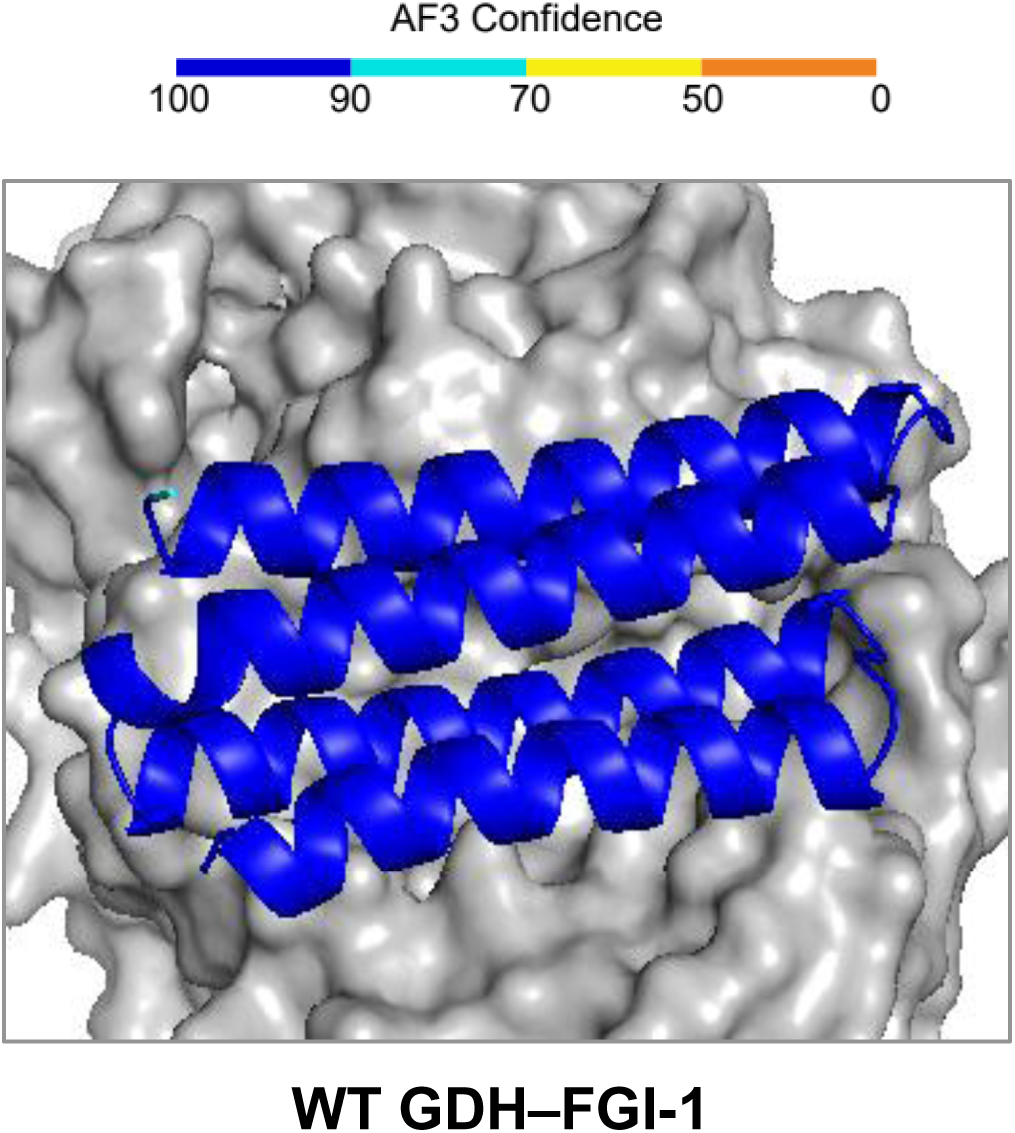
AlphaFold3 prediction of the designed GDH–FGI-1 complex. FGI-1 modeled with WT GDH. FGI-1 is colored according to AlphaFold3 confidence scores, and GDH is shown as a gray surface.

**Figure S2.**
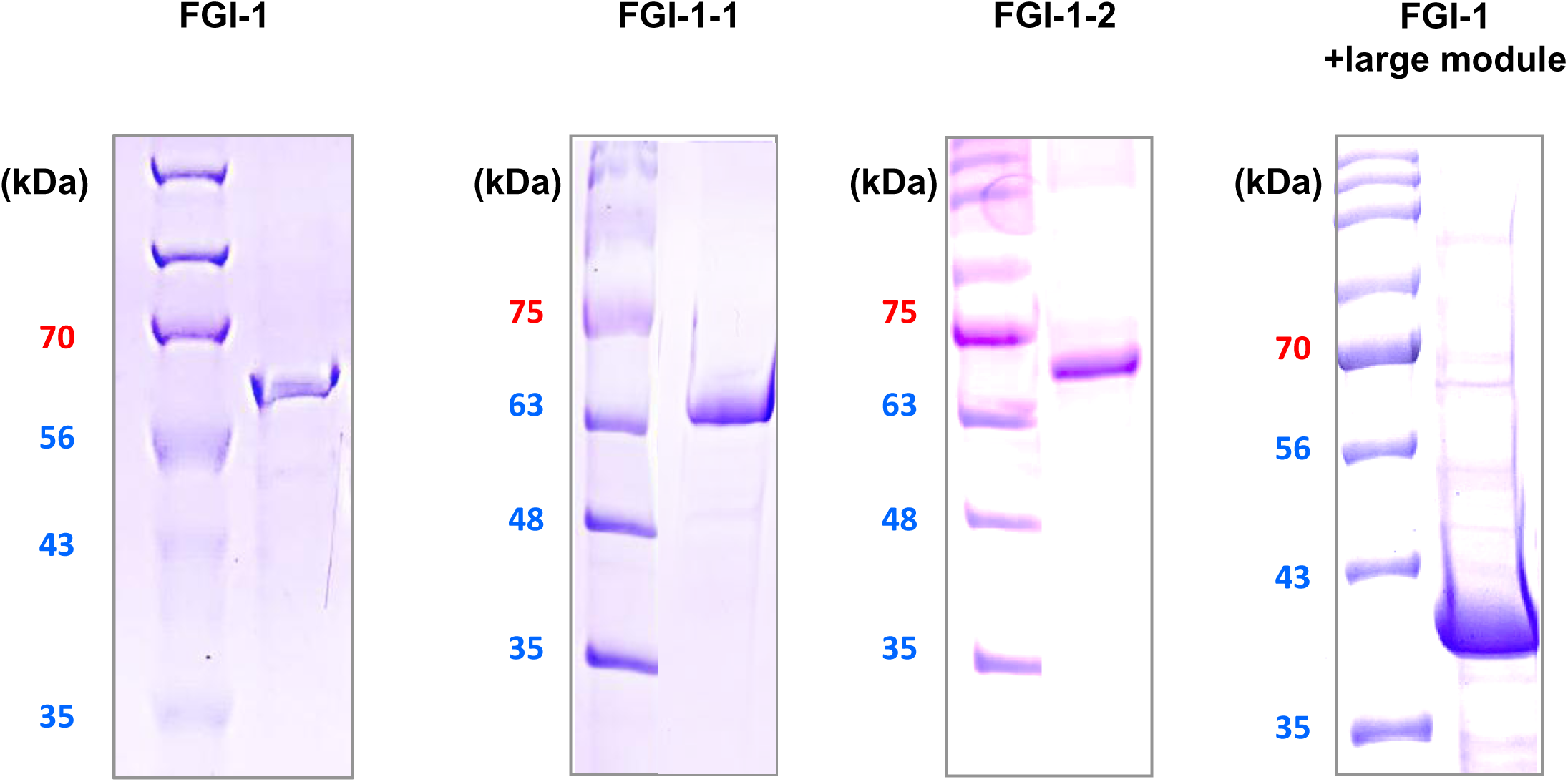
Purification of proteins used for functional assays. Representative SDS–PAGE analysis of purified His–MBP–FGI-1, His–MBP–FGI-1-1, His–MBP–FGI-1-2, and the FGI-1–large-module fusion protein. Molecular-weight markers are indicated in kDa.

**Table S1.** Candidate selection throughout the FGI-1 design and screening workflow. Numbers of candidate sequences retained during Protein MPNN sequence generation, computational filtering, visual prioritization, and FACS-based screening.

| Step | Number of candidates | Description |
| --- | --- | --- |
| Sequence sampling | 500 | ProteinMPNN at 5 sampling temperatures |
| Computational filtering | 88 | Retained after solubility, pLDDT, and interface-quality filtering |
| Visual inspection and prioritization | 6 of 88 | Prioritized by visual inspection for initial FACS screening |
| Initial FACS screening | 6 | Initially prioritized candidates evaluated by FACS |
| FACS screening of remaining candidates | 82 | Subsequently evaluated by FACS |
| Final hit identification | 1 | identified among the 6 initially prioritized candidates |

## Reference

[1] J. Wang, Electrochemical glucose biosensors. Chemical Reviews 2008, 108, 814–825.

[2] S. Tsujimura, K. Kojima, K. Kano, T. Ikeda, M. Sato, H. Sanada, H. Omura, Novel FAD-dependent glucose dehydrogenase for a dioxygen-insensitive glucose biosensor. Bioscience, Biotechnology, and Biochemistry 2006, 70, 654–659.

[3] S. Tsujimura, From fundamentals to applications of bioelectrocatalysis: bioelectrocatalytic reactions of FAD-dependent glucose dehydrogenase and bilirubin oxidase. Bioscience, Biotechnology, and Biochemistry 2019, 83, 39–48.

[4] J. Okuda-Shimazaki, H. Yoshida, K. Sode, FAD dependent glucose dehydrogenases—Discovery and engineering of representative glucose sensing enzymes. Bioelectrochemistry 2020, 132, 107414.

[5] K. Ozawa, H. Iwasa, N. Sasaki, N. Kinoshita, A. Hiratsuka, K. Yokoyama, Identification and characterization of thermostable glucose dehydrogenases from thermophilic filamentous fungi. Applied Microbiology and Biotechnology 2017, 101, 173–183.

[6] S. D. Wijayanti, L. Sützl, A. Duval, D. Haltrich, Characterization of fungal FAD-dependent AA3_2 glucose oxidoreductases from hitherto unexplored phylogenetic clades. Journal of Fungi 2021, 7, 873.

[7] T. Adachi, T. Fujii, M. Honda, Y. Kitazumi, O. Shirai, K. Kano, Direct electron transfer-type bioelectrocatalysis of FAD-dependent glucose dehydrogenase using porous gold electrodes and enzymatically implanted platinum nanoclusters. Bioelectrochemistry 2020, 133, 107457.

[8] A. Navaee, A. Salimi, FAD-based glucose dehydrogenase immobilized on thionine/AuNPs frameworks grafted on amino-CNTs: Development of high power glucose biofuel cell and biosensor. Journal of Electroanalytical Chemistry 2018, 815, 105–113.

[9] G. Sakai, K. Kojima, K. Mori, Y. Oonishi, K. Sode, Stabilization of fungi-derived recombinant FAD-dependent glucose dehydrogenase by introducing a disulfide bond. Biotechnology Letters 2015, 37, 1091–1099.

[10] H. Yoshida, G. Sakai, K. Mori, K. Kojima, S. Kamitori, K. Sode, Structural analysis of fungus-derived FAD glucose dehydrogenase. Scientific Reports 2015, 5, 13498.

[11] T. Kortemme, De novo protein design—from new structures to programmable functions. Cell 2024, 187, 526–544.

[12] A. Winnifrith, C. Outeiral, B. L. Hie, Generative artificial intelligence for de novo protein design. Current Opinion in Structural Biology 2024, 86, 102794.

[13] J. Zhou, M. Huang, Navigating the landscape of enzyme design: from molecular simulations to machine learning. Chemical Society Reviews 2024, 53, 8202–8239.

[14] J. L. Watson, D. Juergens, N. R. Bennett, B. L. Trippe, J. Yim, H. E. Eisenach, W. Ahern, A. J. Borst, R. J. Ragotte, L. F. Milles, B. I. M. Wicky, N. Hanikel, S. J. Pellock, A. Courbet, W. Sheffler, J. Wang, P. Venkatesh, I. Sappington, S. V. Torres, A. Lauko, B. De Bortoli, E. Mathieu, S. Ovchinnikov, R. Barzilay, T. S. Jaakkola, F. DiMaio, M. Baek, D. Baker, De novo design of protein structure and function with RFdiffusion. Nature 2023, 620, 1089–1100.

[15] J. Dauparas, I. Anishchenko, N. Bennett, H. Bai, R. J. Ragotte, L. F. Milles, B. I. M. Wicky, A. Courbet, R. J. de Haas, N. Bethel, P. J. Y. Leung, T. F. Huddy, S. Pellock, D. Tischer, F. Chan, B. Koepnick, H. Nguyen, A. Kang, B. Sankaran, A. K. Bera, N. P. King, D. Baker, Robust deep learning-based protein sequence design using ProteinMPNN. Science 2022, 378, 49–56.

[16] J. Jumper, R. Evans, A. Pritzel, T. Green, M. Figurnov, O. Ronneberger, K. Tunyasuvunakool, R. Bates, A. Žídek, A. Potapenko, A. Bridgland, C. Meyer, et al., Highly accurate protein structure prediction with AlphaFold. Nature 2021, 596, 583–589.

[17] J. Abramson, J. Adler, J. Dunger, R. Evans, T. Green, A. Pritzel, O. Ronneberger, L. Willmore, A. J. Ballard, J. Bambrick, S. W. Bodenstein. et al., Accurate structure prediction of biomolecular interactions with AlphaFold 3. Nature 2024, 630, 493–500.

[18] E. T. Boder, K. D. Wittrup, Yeast surface display for screening combinatorial polypeptide libraries. Nature Biotechnology 1997, 15, 553–557.

[19] F. H. Arnold, Directed evolution: bringing new chemistry to life. Angewandte Chemie International Edition 2018, 57, 4143–4148.

